# Super-resolution expansion microscopy reveals nanoscale protein domains and CO_2_-dependent remodeling of Chlamydomonas pyrenoid-traversing membranes

**DOI:** 10.64898/2026.06.11.731689

**Authors:** Aastha Garde, Haoyu Wu, Martin Jonikas

## Abstract

Within the algal carbon-assimilating organelle, the pyrenoid, specialized traversing membranes perform the essential function of delivering concentrated CO_2_ to Rubisco. In *Chlamydomonas reinhardtii*, these membranes consist of peripheral cylindrical tubules that connect to a central reticulated region. However, due to resolution limitations, the spatial distribution of their structural and functional proteins has remained unclear. Here, we achieve an ∼11-fold improvement in resolution by combining ultrastructure expansion microscopy with super-resolution instantaneous structured illumination microscopy, revealing protein localizations and condition-dependent remodeling of these membranes. At air levels of CO_2_, the tubule-initiating protein SAGA1 forms narrow rings at the pyrenoid edge, the tubule-extending protein MITH1 surrounds the peripheral tubules, and the putative transporter BST4 surrounds tubules more centrally, suggesting that the cylindrical tubules contain multiple distinct protein domains. The CO_2_-delivering carbonic anhydrase CAH3 localizes to the inner face of the central reticulated region, suggesting that this region is specialized for CO_2_ delivery. CAH3 remains in the reticulated region at high CO_2_, suggesting that the cell maintains a minimal CO_2_-delivery apparatus even when dispensable. Finally, at high CO_2_, cylindrical tubule diameter narrows, and MITH1 relocalizes throughout the pyrenoid-traversing membrane network. Together, our study elucidates sub-pyrenoid protein organization and CO_2_-dependent reorganization.

## INTRODUCTION

The pyrenoid, an organelle found in the chloroplasts of many algal species, is responsible for approximately one-third of global photosynthetic CO₂ fixation^1–4^. To date, the pyrenoid of the genetically tractable model alga *Chlamydomonas reinhardtii* (Chlamydomonas hereafter) remains the best molecularly and morphologically characterized^5,6^. It consists of the matrix, a liquid-like condensate^7^ of the CO_2_-fixing enzyme Rubisco clustered by the linker protein EPYC1 (Cre10.g436550)^4,8,9^; a network of thylakoid membrane-derived traversing membranes^10–12^ that deliver concentrated CO_2_ to Rubisco^13–15;^ and a surrounding starch sheath that likely serves as a diffusion barrier to prevent CO_2_ loss (Figure 1a-b)^16,17^. The CO_2_-delivery function of the pyrenoid-traversing membranes is achieved by the thylakoid carbonic anhydrase CAH3 (Cre09.g415700) housed in the lumen^16,18–21^. CAH3 converts abundant luminal HCO_3_^-^to concentrated CO_2_, driven by the low pH environment of the lumen^15,22^. This concentrated CO_2_ then diffuses out of the pyrenoid-traversing membranes to enhance the enzymatic activity of the Rubisco in the matrix. There is considerable interest in engineering algal pyrenoids into crops, as mathematical modeling suggests that this could improve yields and increase both water and nitrogen use efficiency^16,23–25^.

**Figure 1:**
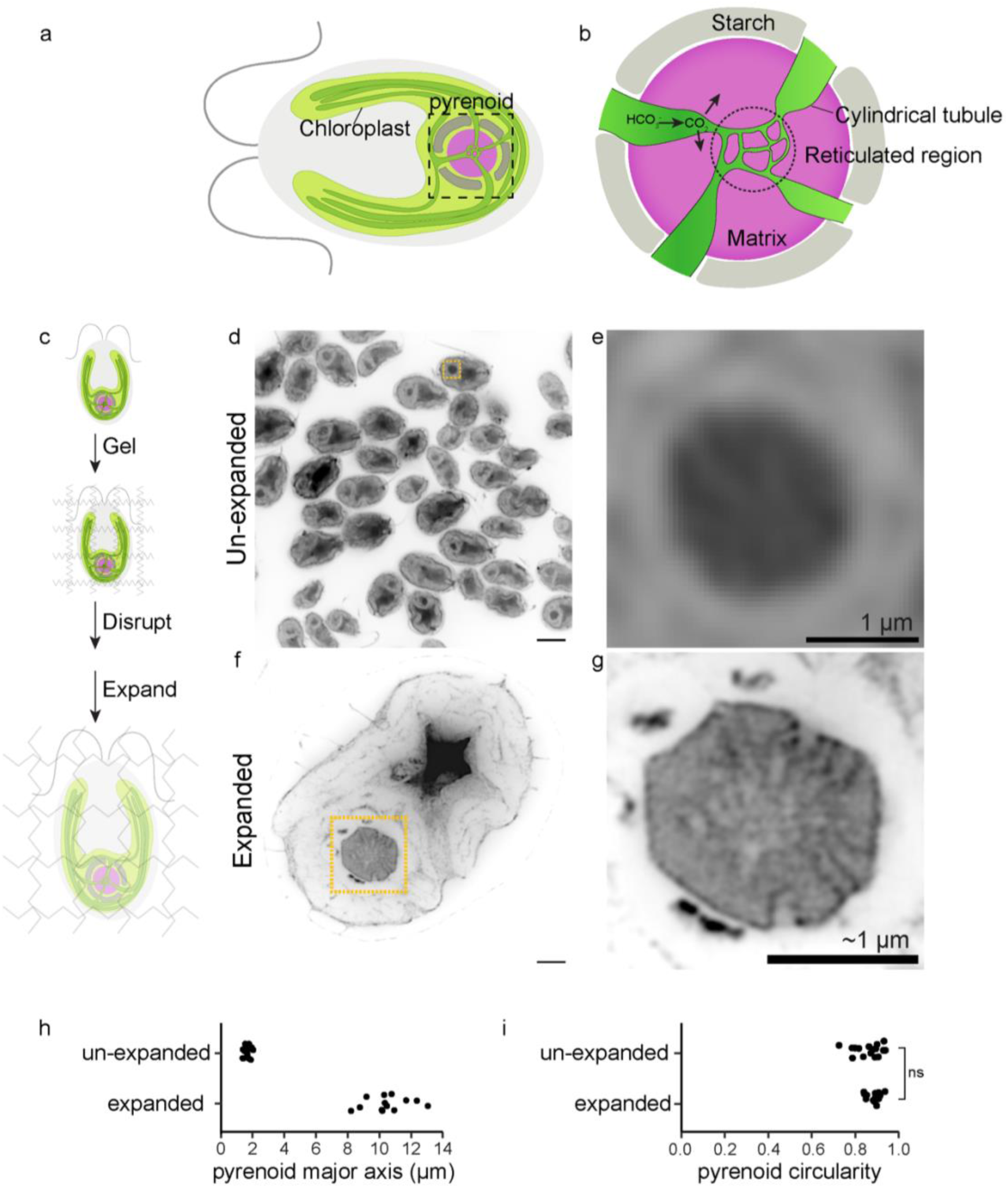
Ultrastructure Expansion Microscopy (U-ExM) reveals detailed pyrenoid architecture. **a.** Cartoon depiction of the Chlamydomonas cell; the pyrenoid is shown at the base of the cup-shaped chloroplast. **b.** Zoomed-in view of a cartoon depiction of the pyrenoid. **c.** Schematic of U-ExM workflow. **d.** Field-of-view of unexpanded Chlamydomonas cells stained with NHS ester pan-protein stain. The pyrenoid is boxed in dotted lines (scale bar 5 µm). **e**. Zoomed-in view of the pyrenoid of an unexpanded cell from **d** (scale bar 1 µm). **f**. Field-of-view of an expanded Chlamydomonas cell stained with pan-protein stain. The pyrenoid is boxed in dotted lines (scale bar 5 µm, not corrected for expansion factor). **g**. Zoomed-in view of the pyrenoid from the expanded cell in **f** (scale bar ∼1 µm, corrected for expansion factor). **h.** Quantification of the major axis of the pyrenoid in un-expanded and expanded cells. **i**. Quantification of pyrenoid circularity in unexpanded and expanded cells. Cells were harvested after growing at low CO_2_ for 6 hours. Statistical p-values were calculated using a two-tailed t-test.

While the pyrenoid matrix has been structurally and functionally well-characterized, understanding the proteins involved in the biogenesis and shaping of the pyrenoid-traversing membranes is the next frontier in pyrenoid biology. At air levels of CO_2_ (0.04% v/v), the pyrenoid-traversing membranes are defined by two distinct morphological domains, cylindrical tubules and a reticulated region, that have been described first by transmission electron microscopy (TEM)^12,26^ and more recently by cryo-electron tomography (cryoET)^10^. Near the periphery of the pyrenoid, flat thylakoid sheets merge into cylindrical tubules ∼100 nm in diameter, which radially enter the matrix. Near the center of the pyrenoid, the cylindrical tubules narrow in diameter, merge with each other via three-way junctions, and form the reticulated region (Figure 1a-b). Why the pyrenoid-traversing membranes exhibit these two distinct morphological domains remains a major open question in the field.

Understanding the potential functions of the pyrenoid-traversing membranes’ domains requires precisely mapping membrane protein localization, as knowledge of a protein’s subdomain localization is critical to understanding its potential function within the pyrenoid-traversing membranes. In addition to the CO_2_-delivering CAH3, several other key players that function in or on the pyrenoid-traversing membranes have been identified, including the putative transmembrane transporter BST4 (Cre06.g261750), the tubule-initiating protein SAGA1 (Cre11.g467712), and the tubule-extending protein MITH1 (Cre06.g259100). However, we currently lack a clear picture of their precise spatial organization. Furthermore, while traversing-membrane morphology has been well-studied at air levels of CO_2_, the pyrenoid is known to be smaller under high CO_2_ conditions^4,27^, and whether traversing-membrane morphology and protein localization undergo remodeling under high CO_2_ conditions also remains uncharacterized.

A major obstacle to mapping the precise spatial organization of key factors has been that the small diameter and tight packing of the traversing membranes make them challenging to resolve by conventional confocal microscopy. Super-resolution light microscopy techniques like Structured Illumination Microscopy (SIM)^28^ and Stimulated Emission Depletion (STED)^29,30^ are a possible avenue; however, both techniques only provide an additional two- to four-fold improvement in resolution (∼60-120 nm lateral resolution) in ideal conditions, which is still insufficient to visualize pyrenoid-traversing membranes and their structural subdomains in detail.

To overcome these technical challenges to visualizing protein localization within the pyrenoid-traversing membranes, here we combine Ultrastructure Expansion Microscopy (U-ExM) with super-resolution instantaneous SIM (iSIM)^31^. U-ExM has been used previously with Chlamydomonas cells to visualize protein localization and cellular morphology, with a focus on the cytoskeleton and flagellar basal bodies^32–34^. We adapt the Chlamydomonas U-ExM protocol to study pyrenoid-traversing membrane protein organization. We show that this method enables clear visualization of the three-dimensional architecture of the wild-type pyrenoid-traversing membranes. By combining immunofluorescence with U-ExM + iSIM, we discover distinct radial localizations of four known pyrenoid membrane proteins, notably suggesting a role for the central reticulated region in localizing the essential CO_2_-delivering carbonic anhydrase CAH3. Finally, we discover that the morphology and protein distribution of the pyrenoid-traversing membranes respond to changes in environmental CO_2_ concentrations. Together, these discoveries advance the understanding of the molecular architecture of the pyrenoid.

## RESULTS

### Ultrastructure Expansion Microscopy with iSIM achieves ∼24 nm resolution of Chlamydomonas pyrenoid morphology

To achieve an effective 10- to 12-fold increase in fluorescence imaging resolution of pyrenoid ultrastructure, we combined an enhanced expansion microscopy protocol with instantaneous structured illumination microscopy (iSIM). In an expansion microscopy workflow, cells are embedded in a swellable hydrogel, the structural molecules are denatured to make the sample mechanically homogenous, and then the sample is physically expanded, which improves the resolution achievable by light microscopy (Figure 1c). We first optimized the existing Chlamydomonas U-ExM workflow^32^ to increase the expansion factor from four-fold to approximately six-fold (see Methods). Next, we imaged the expanded cells with iSIM to gain an additional two-fold increase in lateral and axial resolution^31^. We visualized the global protein distribution of the Chlamydomonas cells using a fluorescently conjugated NHS-ester^35–37^, a compound that reacts with primary amines on proteins. As observed previously in cryo-ExM studies of Chlamydomonas cells^33,34^, the protein-dense pyrenoid appears as a densely stained sphere at its canonical position at the base of the cup-shaped chloroplast in both unexpanded (Figure 1d-e, S1a-b) and expanded cells (Figure 1f-g, S1c-d). By quantifying pyrenoid diameter before and after expansion, we found that our modified gel chemistry yielded an average expansion factor of ∼6-fold (Figure 1h), compared to the ∼4-fold achieved using the original U-ExM protocol. In addition, this enhanced expansion factor did not compromise pyrenoid circularity (Figure 1i). Thus, by combining modified U-ExM gel chemistry with iSIM imaging, we achieved an effective lateral resolution of ∼24 nm without distorting the native shape of the pyrenoid.

### U-ExM enables three-dimensional examination of pyrenoid-traversing membrane morphology

As reported previously in expanded pyrenoids^34^, we observed sparsely-stained structures arranged radially within a densely-stained background (Figure 1g). It had been previously hypothesized that the sparsely-stained structures were the pyrenoid-traversing membranes, and the densely-stained background was the matrix^34^, but these hypotheses had not been experimentally tested.

To examine how these protein-sparse structures were organized across the whole organelle, we imaged the total volume of pyrenoids from wild-type cells grown photoautotrophically at air levels of CO_2_ for six hours before fixation. In superficial slices of wild-type pyrenoids, we observed that the densely stained pyrenoid matrix was punctuated with 3-5 circular protein-sparse shapes, which we hypothesized were cross-sections of cylindrical tubules entering the matrix. In medial slices of wild-type pyrenoids, we instead observed protein-sparse tubular shapes that met in the middle of the pyrenoid in a complex geometry, which we hypothesized were longitudinal sections of the cylindrical tubules meeting in a reticulated network (Figure 2a, S2a, Movie S1).

**Figure 2:**
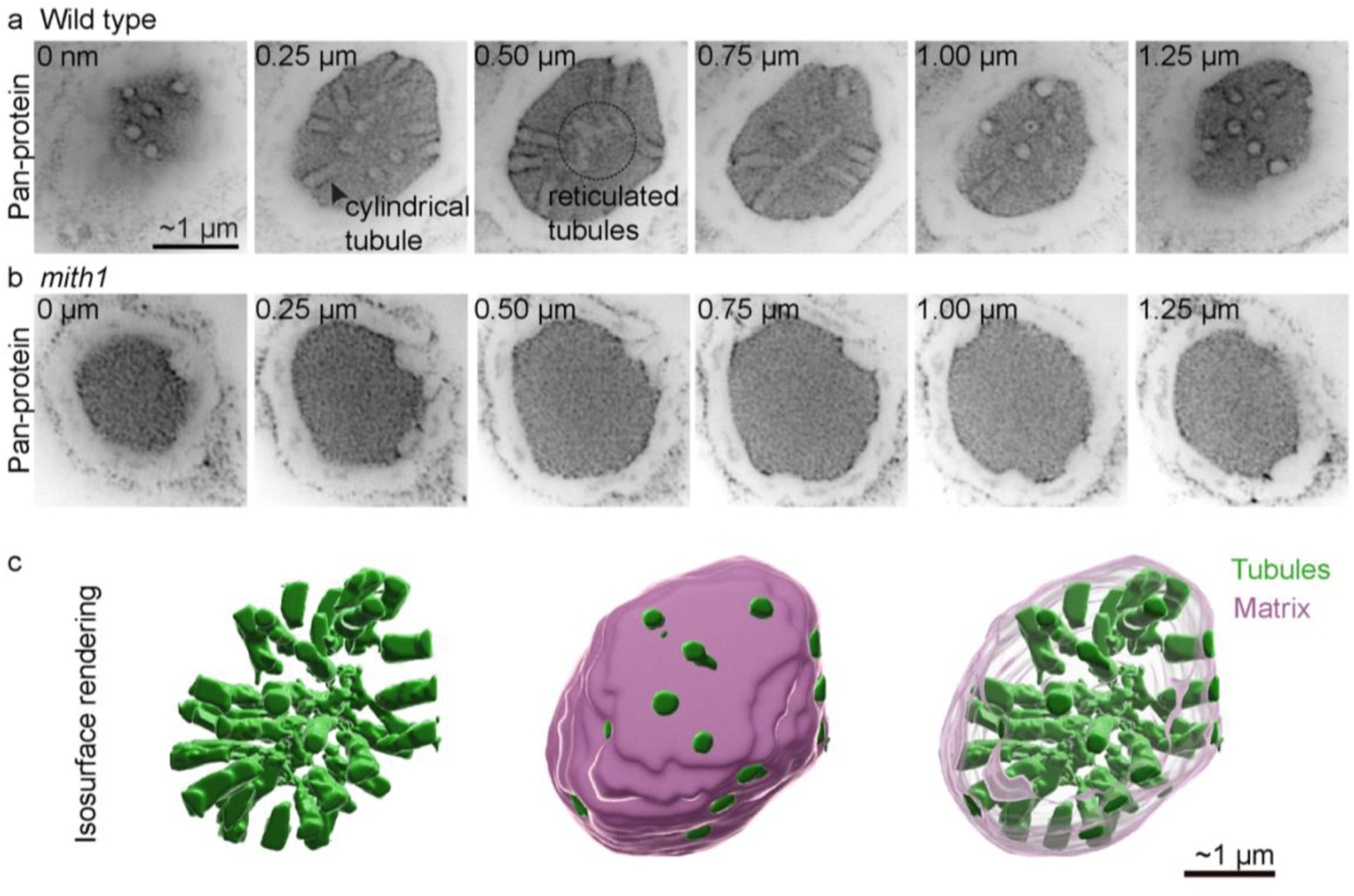
U-ExM enables three-dimensional characterization of pyrenoid-traversing membrane morphology. **a.** Confocal z-sections of a wild-type pyrenoid stained with pan-protein stain. Expansion corrected relative z-position depicted in the top left corner. **b.** Confocal sections of a *mith1* mutant pyrenoid stained with pan-protein stain. Expansion corrected relative z-position depicted in the top left corner. **c.** Space-filling reconstructions of tubules and matrix segmented from the confocal sections in **a**. Cells were harvested after growing at low CO_2_ for 6 hours. Scale bars were corrected for expansion factor.

To test the hypothesis that the protein-sparse structures are indeed pyrenoid-traversing membranes, we expanded and compared the morphology of wild-type pyrenoids to those of the *mith1* mutant. Because the *mith1* mutant lacks the tubule extension protein MITH1, it lacks traversing membranes within the pyrenoid matrix^29^. Consistent with the above hypotheses, *mith1* mutants had a clear, large, spherical, protein-dense matrix at the base of the chloroplast but lacked the protein-sparse structures within the matrix (Figure 2b, S2b). Comparing this known mutant phenotype to wild-type cells gave us confidence that the protein-dense and protein-sparse pan-protein staining compartments we observed in wild-type cells by U-ExM correspond to the pyrenoid matrix and pyrenoid-traversing membranes, respectively.

Furthermore, in wild-type pyrenoids, we noticed an increase in pan-protein staining density at the edges of the cylindrical tubules (Figure 2a, arrowhead) that was absent from the more central reticulated region. This staining density might correspond to a unique protein domain on or adjacent to the cylindrical tubule membrane.

By sampling in the Z-dimension every 200 nm (corresponding to ∼33 nm pre-expansion), we could use our confidence in the structures we observed by pan-protein staining to segment and reconstruct the entire pyrenoid-traversing membranes of wild-type cells (Figure 2c, S2c-d, Movie S2). We observed that wild-type pyrenoids had cylindrical tubules entering radially throughout the matrix. Consistent with prior cryo-ET studies of the pyrenoid^10^, these tubules narrowed in diameter as they approached the center of the pyrenoid and interconnected to form a network of reticulated tubules^38^.

Prior cryo-ET imaging had also shown pyrenoid tubules entering adjacent to starch plates in individual lamellae^10^. Our expanded 3D reconstructions allowed us to examine these tubule entry sites across a larger region of the pyrenoid surface. While we cannot directly visualize the starch plates by U-ExM, we found that tubule entry sites were not randomly distributed around the surface of the pyrenoid; instead, multiple tubules entered the pyrenoid in linear arrangements that we speculate correspond to the boundaries of starch plates (Figure S2e). Together, our observations establish that U-ExM can enable the characterization of pyrenoid-traversing membrane morphology under different growth conditions and in mutants.

### The pyrenoid-traversing membrane proteins MITH1 and BST4 localize to spatially distinct subdomains

Existing light microscopy studies of pyrenoid proteins have been able to distinguish between a protein localized to the matrix, membranes, and starch sheath, but have had very limited ability to determine the localization of a protein within a subdomain of the pyrenoid-traversing membranes^5,6,29,38,39^. By combining immunofluorescence with NHS-ester based pan-protein staining, we determined the distribution of two known pyrenoid tubule-localized proteins with highly specific antibodies: the tubule-extending protein MITH1^29^ and the putative transmembrane transporter BST4 (also called RBMP1)^39^. Previously, both MITH1 and BST4 have been localized to the pyrenoid-traversing membranes via their colocalization with the chlorophyll-positive thylakoid signal in the pyrenoid^29,39^. We reasoned that any potential subdomain-specific localization within the network could reflect functional specialization of the proteins, the subdomain, or both.

Given the approximately circular cross-section of the pyrenoid, we quantitatively compared the radial distributions of MITH1 and BST4 as a function of the distance from the centroid of the pyrenoid. We found that while the pan-protein staining was evenly distributed across the axis of the pyrenoid, MITH1 signal was largely restricted to the cylindrical tubules at the pyrenoid periphery (Figure 3a-b, S3a, Movie S3). The distribution of the transmembrane BST4^38,39^ occupied an intermediate radial distribution, being less enriched in the reticulated region at the center of the pyrenoid and not extending as far peripherally as MITH1 (Figure 3e-f, S3b, Movie S3). Our previous work has shown that the *mith1* mutant fails to extend tubules through the pyrenoid matrix^29^. Our current observation that MITH1 is enriched on the peripheral tubules suggests that MITH1 may be involved in initiating and maintaining the outermost tubules, while the central tubules may be extended and shaped by additional uncharacterized proteins.

**Figure 3:**
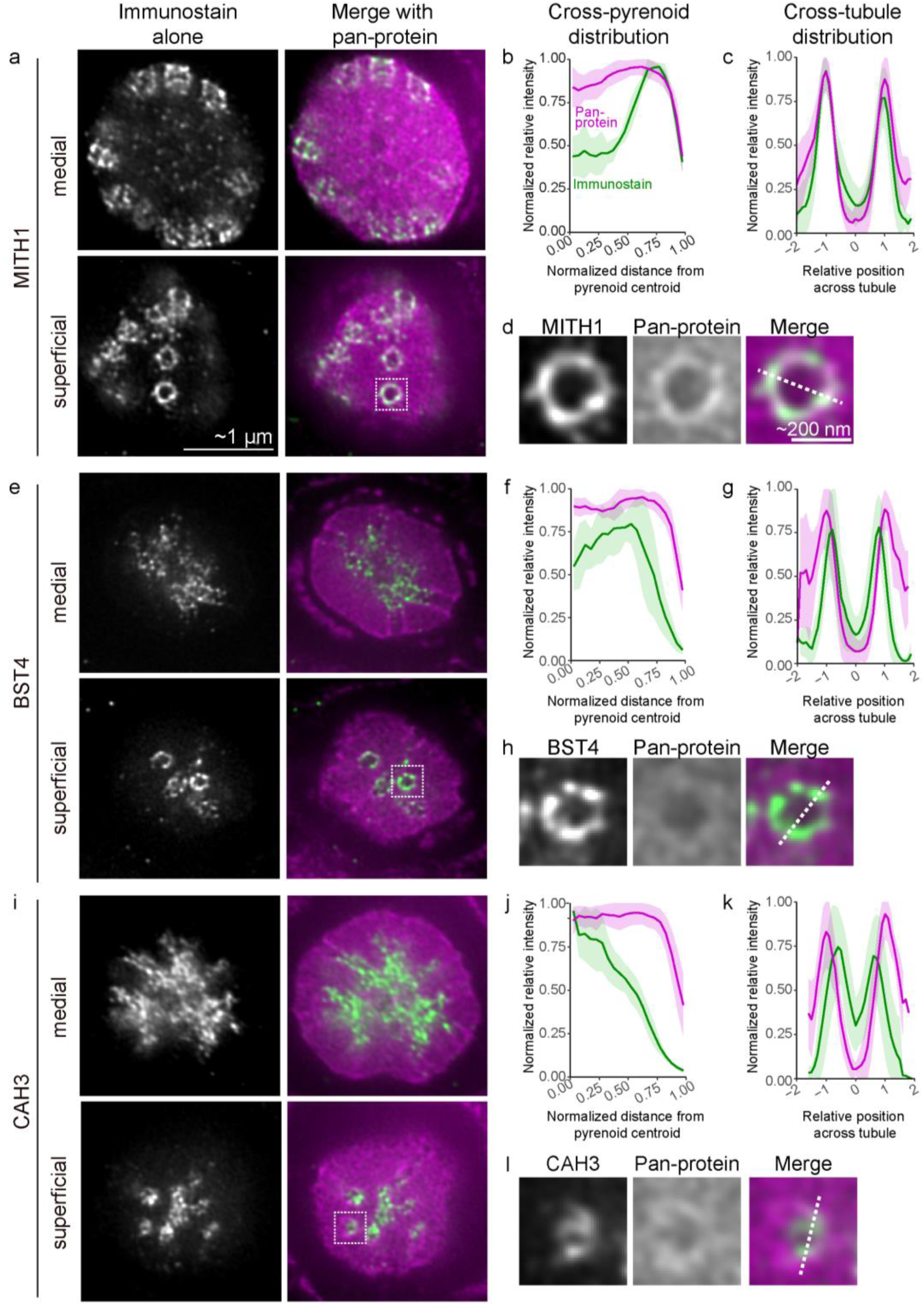
Pyrenoid-traversing membrane proteins are organized in subdomains. **The antibodies used are as follows: a-d.** MITH1, **e-h.** BST4 and **i-l.** CAH3. Wild-type pyrenoids displayed with immunofluorescence alone (gray) and immunofluorescence (green) merged with pan-protein stain (magenta). Medial and superficial slices are displayed for each pyrenoid. **b, f, j.** Cross-pyrenoid distribution: plot profiles of the signal intensity of immunofluorescence (green) and pan-protein stain (magenta) as a function of distance from the centroid of the pyrenoid. The x-axis displays a normalized distance from centroid where the maximum pyrenoid diameter is set to 1. (N ≥5 pyrenoids per condition) **c, g, k.** Cross-tubule distribution: line scan profiles across the tubule axis of the signal intensity of immunofluorescence (green) and pan-protein stain (magenta) as a function of relative distance from the center of the tubule lumen. X-axis displays normalized positional units across the tubules such that the trough in pan-protein stain across multiple tubules could be aligned at 0. (N ≥10 tubules per condition). **d, h, l:** Zoomed-in views of the boxed region of superficial slices displaying individual tubules with immunofluorescence alone, pan-protein stain alone and merged images. Dotted lines depict representative line scans across the tubule axis. Cells were harvested after growing at low CO_2_ for 6 hours. Scale bars were corrected for expansion factor. All plots depict the mean distribution with the shaded region representing the standard deviation from the mean.

Next, we examined protein localization across the axis of individual tubules using the tubule edge-associated increased pan-protein staining density as a fiducial marker. On the peripheral tubules, we found that MITH1 signal overlapped with the increased pan-protein stain density (Figure 3c-d), while the transmembrane BST4 signal localized internally relative to the peaks of pan-protein stain. Using transmembrane BST4 as a proxy for the tubule membrane, our results suggest that the antibody epitope-containing C-terminus of MITH1 is localized distally from the tubule membrane (Figure 3g-h). This finding is supported both by our prior live-cell imaging, which similarly observed MITH1-Venus occupying a broader region surrounding the tubules than BST4-Venus^29^ and by recent structural evidence suggesting that MITH1 forms an elongated coiled-coil that extends away from the membrane and into the matrix^40^.

Together, combining immunofluorescence with U-ExM + iSIM revealed that MITH1 is positioned peripheral to BST4 at both the whole-pyrenoid and sub-tubular scales, demonstrating the utility of this approach for resolving spatial protein organization in the pyrenoid. The distinct localizations of MITH1 and BST4 also suggest that the cylindrical tubules possess at least two distinct protein subdomains that are not morphologically apparent by electron microscopy.

### The luminal carbonic anhydrase CAH3 localizes primarily to the reticulated region

The thylakoid carbonic anhydrase CAH3 is essential for pyrenoid function^14,18,20,21^ and is thought to facilitate the release of concentrated CO_2_ from the pyrenoid tubules by converting HCO_3_^-^ to CO_2_^15,22^. Prior immunogold EM with anti-CAH3 antibodies observed immunogold particles associated largely with the pyrenoid tubules, with fewer particles associated with the surrounding thylakoid sheets^41–43^. More recent immunofluorescence-based localization observed CAH3 to be localized almost exclusively to the pyrenoid tubules^29^. However, the specific localization of CAH3 within the tubule network has remained unexplored due to limitations in the sensitivity and spatial resolution of the existing techniques.

We used immunofluorescence in combination with U-ExM + iSIM to examine the localization of CAH3 in the pyrenoid. Like the prior immunodetection studies, we also observed the vast majority of CAH3 signal in the pyrenoid tubules. However, in contrast to the localization of MITH1 and BST4, we observed that CAH3 signal was localized almost exclusively near the centroid of the pyrenoid in the region corresponding to the narrower tubules of the reticulated region (Figure 3i-j, S3c, Movie S3). These results indicate that CAH3 localizes primarily to the reticulated region of the pyrenoid-traversing membranes.

### CAH3 is on the tubule membrane, not soluble in the lumen

Prior biochemical experiments had found that CAH3 co-fractionates with PSII-rich thylakoid membranes^18,41,44^, has a twin-arginine thylakoid lumen targeting sequence^18^, and is resistant to protease treatment^18,45^, suggesting that it localizes to the lumen of the pyrenoid-traversing membranes. However, it had remained unclear whether CAH3 is soluble, or membrane-associated within the lumen. When we quantified the distribution of CAH3 across the axis of individual tubules, we observed that CAH3 formed two distinct peaks at the edges of the tubules, indicating that CAH3 is not soluble, but rather is closely associated with the membranes. The CAH3 signal localized further inside the pan-protein peaks than the peaks of BST4 (Figure 3k-l). These results indicate that CAH3 localizes to the luminal face of the pyrenoid-traversing membranes rather than being soluble within the tubule lumen.

### SAGA1 forms narrow rings around the cylindrical tubules at the surface of the matrix

Generating specific antibodies for every pyrenoid tubule protein is not always feasible. Thus, we leveraged the existing Venus-3xFLAG-tagged fusion protein collection in Chlamydomonas^5,6^ to demonstrate its use in conjunction with U-ExM + iSIM for high-resolution protein localization. We chose to examine the localization of SAGA1-Venus-3xFLAG as it is both essential for tubule biogenesis^29,46^ and does not have a specific antibody^38,46^.

Based on previous studies that localized SAGA1 to puncta at the pyrenoid periphery^29,46,47^, we expected SAGA1 to localize to a peripheral subdomain of the pyrenoid-traversing membranes. Surprisingly, our anti-FLAG immunofluorescence revealed that SAGA1 localized to a very narrow ring at the pyrenoid periphery (Figure 4a-b, S3d). We further confirmed by two-color immunofluorescence of U-ExM samples that SAGA1 and MITH1 occupied distinct radial zones within the pyrenoid (Figure 4e-f, S3e, Movie S4), with SAGA1 being more peripheral than MITH1, adding a fourth layer of radial organization to the pyrenoid-traversing membranes (Figure 4g).

**Figure 4:**
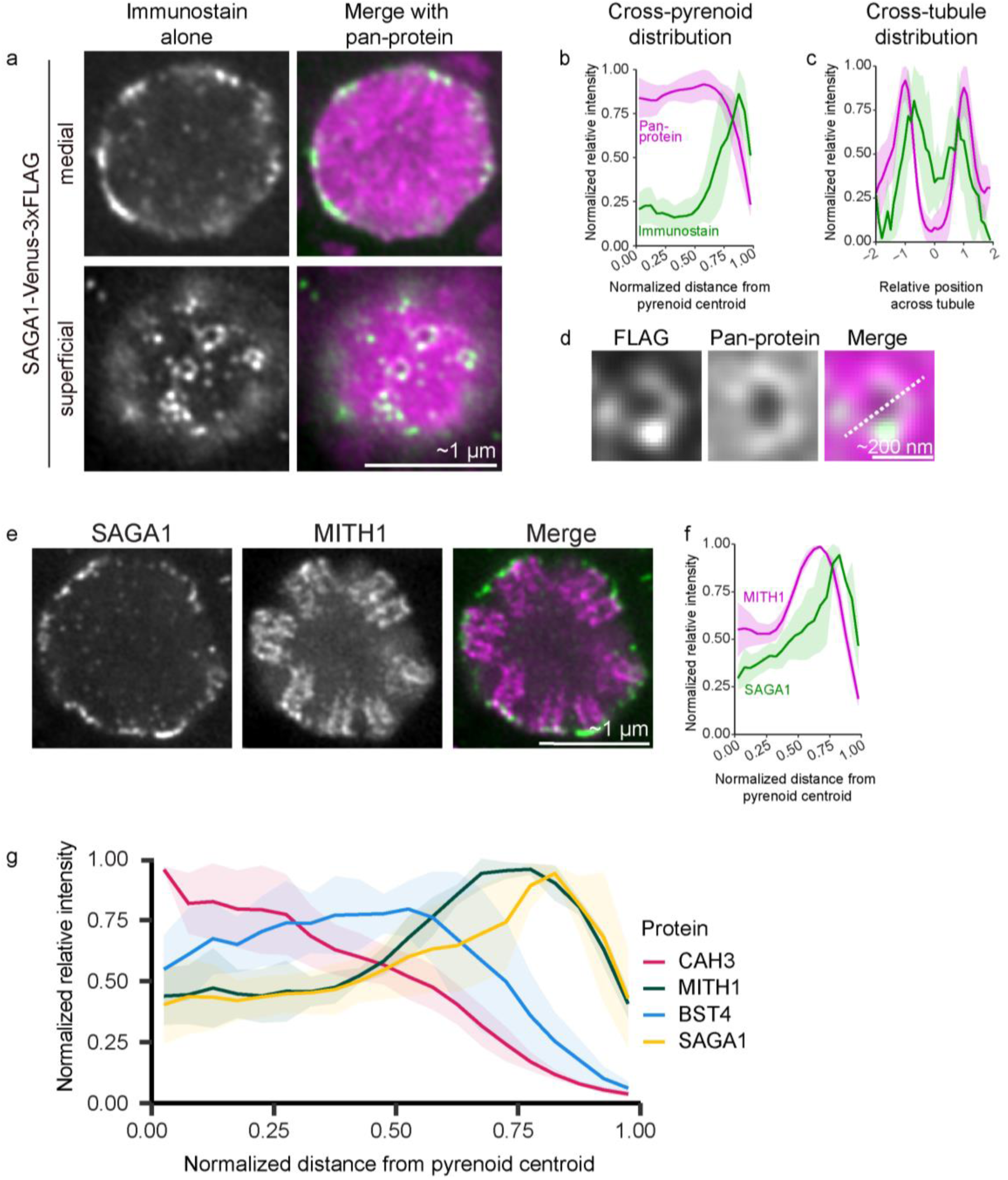
SAGA1 forms rings around the cylindrical tubules at the surface of the matrix. **a.** Pyrenoid of a cell expressing SAGA1-Venus-3xFLAG, displayed with anti-FLAG immunofluorescence alone (gray), and immunofluorescence (green) merged with pan-protein stain (magenta). Both medial and superficial slices are displayed. **b.** Cross-pyrenoid distribution: plot profiles of the signal intensity of immunofluorescence (green) and pan-protein stain (magenta) as a function of distance from the centroid of the pyrenoid (N=6). The x-axis displays a normalized distance from centroid where the maximum pyrenoid diameter is set to 1, shaded areas depict standard deviation from the mean line. **c.** Cross-tubule distribution: line scan profiles across the tubule axis of the signal intensity of immunofluorescence (green) and pan-protein stain (magenta) as a function of relative distance from the center of the tubule lumen. X-axis displays normalized positional units across the tubules such that the trough in pan-protein stain between tubules could be aligned at 0 (N = 10). **d.** Zoomed-in views of the superficial slice of an individual tubule depicting immunofluorescence alone, pan-protein stain alone and merged images. The dotted line depicts a representative line scan across the tubule axis. **e.** Pyrenoid of a wild-type cell expressing SAGA1-Venus-3xFLAG, displayed with FLAG immunofluorescence alone, and MITH1 immunofluorescence alone, and a FLAG (green) and MITH1 (magenta) merged image. **f.** Cross-pyrenoid distribution: plot profiles of the signal intensity of SAGA1 (FLAG, green) and MITH1 (magenta) as a function of distance from the centroid of the pyrenoid (N=6). The x-axis displays a normalized distance from centroid where the maximum pyrenoid diameter is set to 1. Scale bars were corrected for expansion factor. **g.** Combined cross-pyrenoid distribution plot displaying the localization of SAGA1-Venus-3xFLAG (yellow), MITH1 (green), BST4 (blue) and CAH3 (magenta) as a function of distance from the centroid of the pyrenoid. All plots depict the mean distribution with the shaded region representing the standard deviation from the mean. Cells were harvested after growing at low CO_2_ for 6 hours.

Corroborating the evidence that SAGA1 colocalizes with cylindrical tubules, rings of SAGA1 signal co-occurred with protein-sparse tubule entry sites in superficial slices of the pyrenoid (Figure 4a). Across the axis of the tubule, the peaks of SAGA1 signal were more medial than the peak of the pan-protein staining density (Figure 4c-d).

The highly peripheral ring-like localization of SAGA1 at the entry sites of the cylindrical tubules suggests that SAGA1 plays a specialized role at the interface of the pyrenoid-traversing tubules, the pyrenoid matrix, and the chloroplast stroma. SAGA1’s distinct localization compared to MITH1 supports our prior hypotheses^29^ that SAGA1 likely acts first at the surface of the pyrenoid to initiate pyrenoid-traversing membranes’ entrance into the matrix, and then MITH1 acts to extend them through the pyrenoid.

### Tubule diameter increases in response to low CO_2_ concentrations

The pyrenoid is a dynamic organelle whose morphology is known to respond to environmental CO_2_ concentrations. Prior work has documented that after shifting Chlamydomonas cells from growth in high to low CO_2_, the pyrenoid matrix condensate grows^4,27^ and the starch sheath grows to surround the pyrenoid^48^. However, how the morphology of the pyrenoid-traversing membranes changes in response to a transition from high to low CO_2_ has, to our knowledge, not been described.

To study changes in tubule morphology, we cultured wild-type cells at high CO_2_ conditions (HC; 3% CO_2_ v/v) and transferred them to grow under low CO_2_ conditions (air level, 0.04% CO_2_ v/v) for 1 (LC 1h), 3 (LC 3h) and 16 (LC 16h) hours before fixation and preparation for the U-ExM workflow. As others have observed previously^4,49,50^, we saw that HC cells had the smallest pyrenoid matrix diameter, and that pyrenoid diameter increased steadily with time at low CO_2_ (Figure 5a-b).

**Figure 5:**
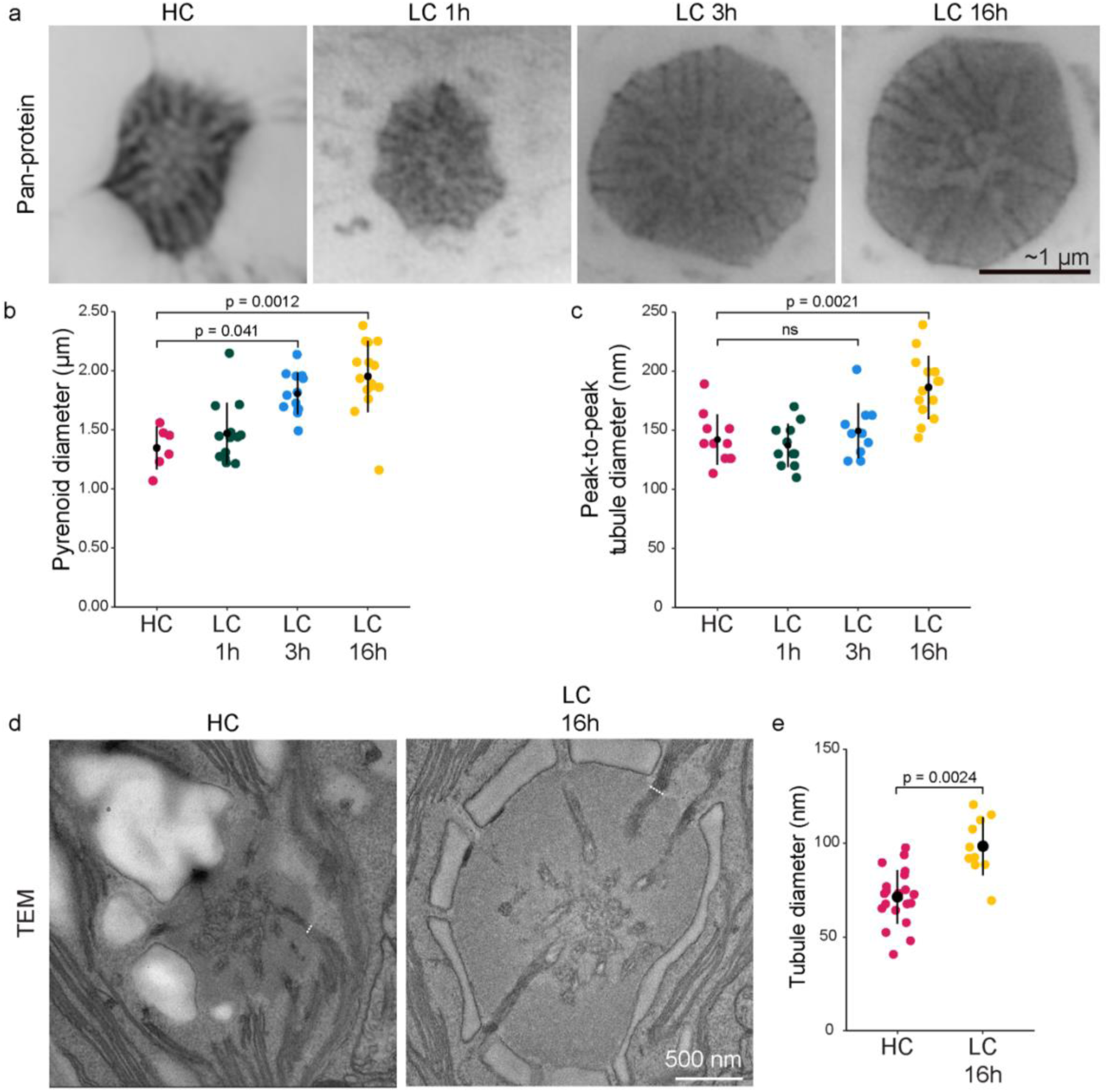
Pyrenoid tubule diameter increases during the transition from high to low CO_2_. **a.** Medial slices of ExM images of wild-type pyrenoids stained with pan-protein stain from cells grown at high CO_2_ (HC) and 1, 3 and 16 hours after transition from high CO_2_ to air levels of CO_2_ (LC 1h, LC 3h, LC 16h). Scale bar was corrected for expansion factor. **b.** Quantification of pyrenoid diameter in each growth condition from ExM images. Error bars depict standard deviation. **c.** Quantification of tubule diameter in each growth condition from ExM images. Black dot depicts mean and error bars depict standard deviation. **d.** Transmission electron micrographs (TEM) of wild-type pyrenoids from cells grown at HC and LC 16h. Dotted lines depict examples of measurements taken for tubule diameter. Scale bar 500 nm. **e.** Quantification of tubule diameter in cells grown at HC and LC 16h from TEMs. Black dot depicts mean and error bars depict standard deviation. Statistical p-values in **b** and **c** were calculated using a Kruskal-Wallis rank sum test followed by a post-hoc Dunn’s test with a Bonferroni correction for multiple comparisons. Statistical p-values in **e** were calculated using a two-tailed t-test.

Interestingly, we observed that cylindrical tubules were narrower in cells cultured at HC, and that tubule diameter increased over time after transition from high to low CO_2_ (Figure 5a, c). While pyrenoid matrix diameter increased after only 1 hour at low CO_2_, we found that the cylindrical tubule diameter changed more slowly, with no visible changes after 1 hour, only slight changes compared to high CO_2_ after 3 hours, and ∼35% wider tubules after 16 hours.

We considered that the changes in tubule diameter could be due to artifacts of the U-ExM sample processing workflow, so we validated our U-ExM results by transmission electron microscopy (TEM). We compared the pyrenoids of cells cultured at high CO_2_ to those cultured at low CO_2_ for 16 hours. Corroborating our findings by U-ExM, cylindrical tubule diameter in cells grown at low CO_2_ for 16 hours was ∼30% wider than that of those grown at high CO_2_ (Figure 5d-e).

Our observations establish that, along with changes in the pyrenoid matrix diameter and starch sheath morphology, the morphology of the pyrenoid-traversing membrane tubules also remodels in response to environmental CO_2_ concentrations, with the timescale of this response being slower than that of the pyrenoid matrix.

### CAH3 remains in the reticulated region in high CO_2_ conditions

We next sought to examine how the localization of CAH3 changed within the pyrenoid-traversing membranes during the transition from high to low CO_2_ conditions. Prior work has shown that CAH3 levels remain unchanged between high and low CO_2_ growth conditions^18,41^, and immunogold electron microscopy studies had suggested that some CAH3 remained localized to the pyrenoid even in high CO_2_ conditions^41^. In our experiments, we observed no change in the localization of CAH3 under high CO_2_ compared to low CO_2_ (Figure 6a). In cells cultured at high CO_2_, CAH3 remained enriched near the center of the pyrenoid. As pyrenoid diameter increased with time at low CO2, the localization of CAH3 remained unchanged, staying enriched in the middle ∼30% of the pyrenoid radius. Similar to CAH3, we observed no change in the cross-pyrenoid localization of BST4 in the transition from high to low CO_2_ (Figure 6b).

**Figure 6:**
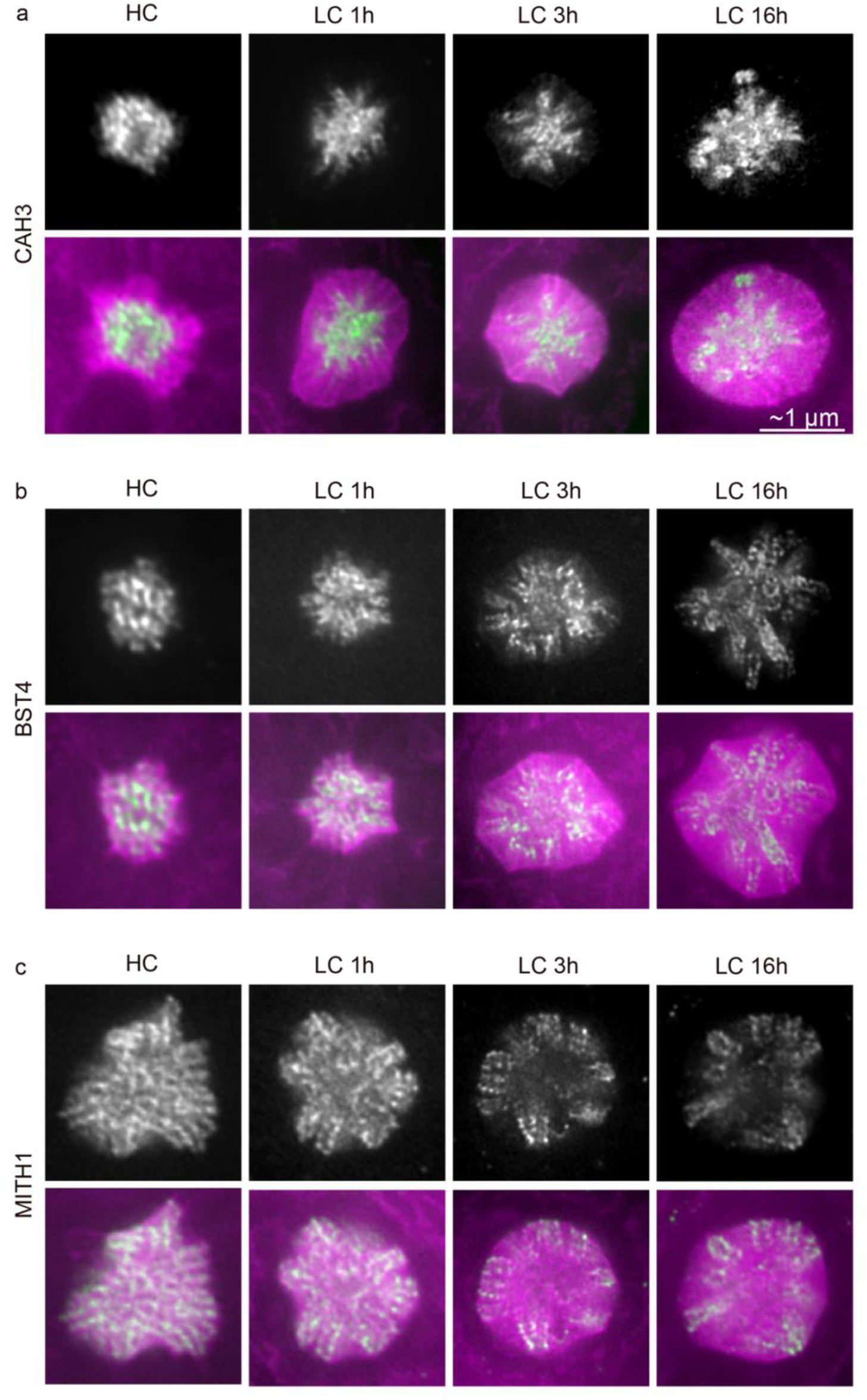
During the transition from high to low CO_2_, MITH1 becomes enriched in the pyrenoid periphery. Medial slices of wild-type pyrenoids displayed with immunofluorescence alone (gray) and immunofluorescence (green) merged with pan-protein stain (magenta) from HC, LC 1h, LC 3h and LC 16h growth conditions. The antibodies used are a. CAH3, b. BST4, and c. MITH1. Scale bars corrected for expansion factor.

The persistent localization of CAH3 and BST4 near the center of the pyrenoid suggests that even at high CO_2_, when the pyrenoid is not required for growth, the cell maintains a minimal CO_2_-delivery apparatus of pyrenoid-traversing membranes, properly localized carbonic anhydrase, and other factors.

### MITH1 relocalizes in response to CO_2_ availability

We next examined the distribution of MITH1 in the pyrenoids of cells grown at high CO_2_ and transitioned to low CO_2_. Unlike CAH3 and BST4, at high CO_2_, MITH1 signal was found throughout the pyrenoid-traversing membranes, including near the center of the pyrenoid. With increasing time at low CO_2_, MITH1 was depleted from the central tubules and adopted its strictly peripheral localization described earlier (Figure 6c). Our results establish that MITH1 localization changes during the transition from high to low CO_2_, potentially in response to CO_2_-induced changes in pyrenoid tubule morphology.

## DISCUSSION

Leveraging the high-resolution imaging provided by U-ExM + iSIM and protein localizations by immunofluorescence allowed us to contribute new insights into sub-pyrenoid protein localizations and how they change from high to low CO_2_ growth conditions (Figure 7).

**Figure 7:**
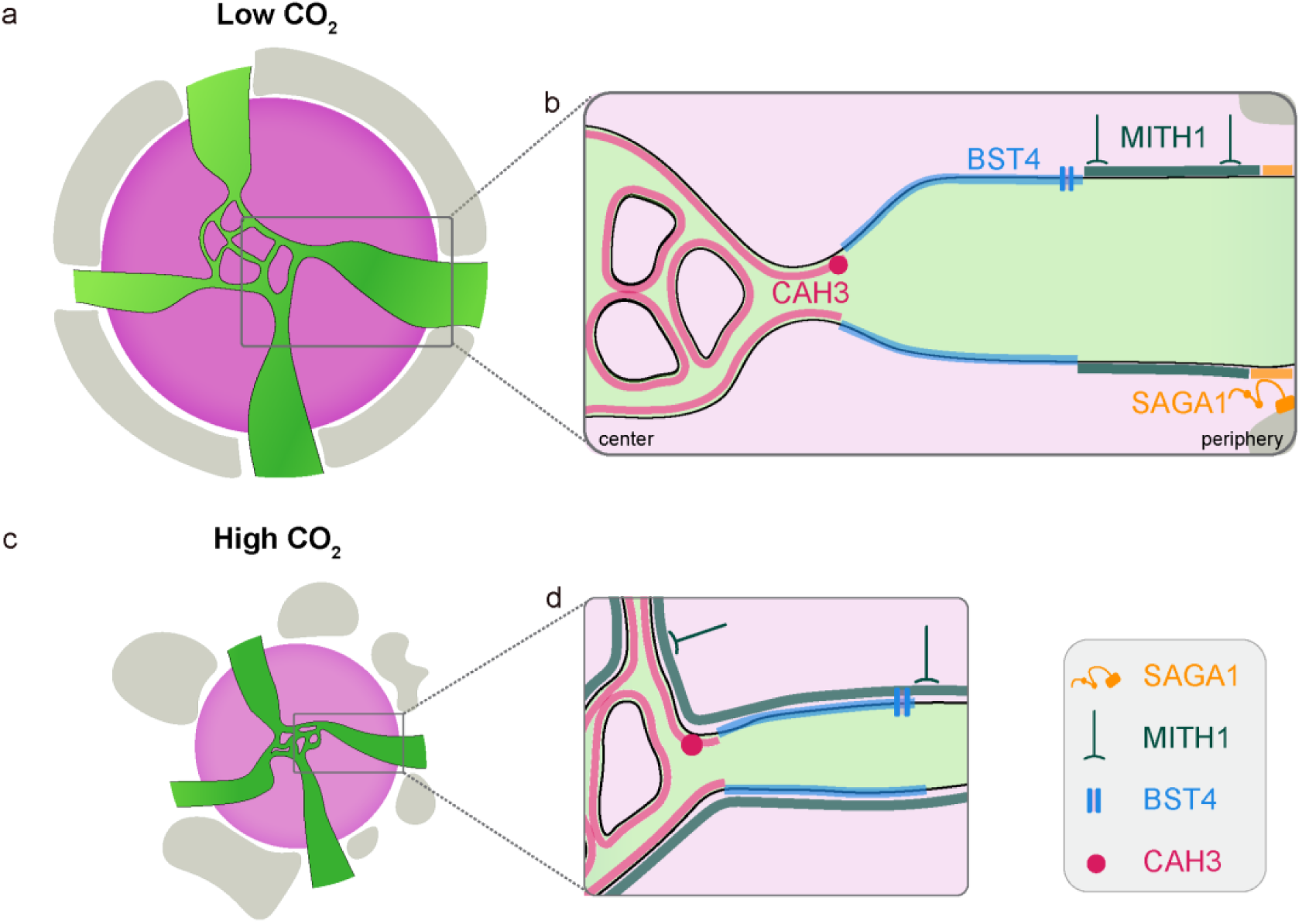
Summary model of our U-ExM-based advances in pyrenoid tubule protein organization. **a.** Cartoon depiction of the pyrenoid in low CO_2_ culture conditions. **b.** Illustration of the cross-pyrenoid and cross-tubule organization of the tubules depicting the relative positions of known tubule proteins in low CO_2_ conditions. **c.** Cartoon depiction of the pyrenoid in high CO_2_ culture conditions depicting narrower cylindrical tubules. **d.** Illustration of the cross-pyrenoid and cross-tubule organization of the tubules depicting the relative positions of known tubule proteins in high CO_2_ conditions.

We found that the key carbonic anhydrase CAH3 localizes specifically to the central reticulated region, suggesting that the reticulated region is specialized for CO_2_ release. If the reticulated region is responsible for CO_2_ release, this suggests that the cylindrical tubules that connect the reticulated region to the surrounding thylakoids may serve as conduits for transport of ions, substrates, and products.

Our findings suggest that CAH3 localizes to the surface of the tubule membrane; however, the mechanistic basis of CAH3’s association with the membrane is unknown. One possibility is that CAH3 can independently associate with membranes as the CAH3 dimer has a predicted hydrophobic region that has been hypothesized to allow it to interact with membranes^51,52^; however, this has not been tested. Alternatively, CAH3 could interact with an unknown membrane-localized targeting factor that localizes CAH3 to the membrane.

Given that the lumen of the pyrenoid tubule network is continuous with that of the chloroplast thylakoids^10^, it is also currently unknown how CAH3 remains restricted to the pyrenoid tubules under all growth conditions. CAH3, independently or via a binding partner, might sense the unique morphology or curvature of the reticulated tubules and preferentially localize to them. Phosphorylation of CAH3 in response to low CO_2_ conditions has also been proposed as a possible mechanism for its localization to the pyrenoid^41^ and could be a regulatory modification used to localize CAH3, either independently or in concert with a binding partner. The installation of CO_2_-delivery machinery is central to engineering functional pyrenoids into land plants; thus, understanding the mechanism of CAH3 localization is a crucial step in the engineering process.

While luminal CAH3 and transmembrane BST4 retained their radial localizations independent of CO_2_ concentrations, we were surprised to find that the radial localization of MITH1 changed from high to low CO_2_. MITH1 was uniformly distributed throughout the tubule network at high CO_2_ and became increasingly peripheral during the transition to low CO_2_. We recently showed that MITH1 can bind both membranes and Rubisco, likely serving to link the pyrenoid tubule and matrix compartments^40^. As the proportion of pyrenoid-localized Rubisco decreases at high CO_2_^4^, the spatial distribution of MITH1 may be driven by the availability and radial extent of Rubisco. Thus, at high CO_2_, MITH1 may ingress deeper into the pyrenoid-traversing tubule network to prevent unsatisfied Rubisco-binding sites and maintain connection with both membranes and matrix.

Previous electron microscopy-based studies had documented the persistence of the pyrenoid-traversing membranes in cells cultured at high CO_2_^48^. We found that the cylindrical tubule diameter is smaller in cells grown at high CO_2_ than in those grown at low CO_2_. We additionally found that the increase in tubule diameter lags behind the increase in matrix diameter during the high-to-low CO_2_ growth transition. These observations suggest that the pyrenoid-traversing membranes are slower to remodel in response to changing CO_2_ conditions and might explain why they persist in growth conditions where the function of the pyrenoid is not required.

Existing U-ExM workflows are increasingly being used and adapted to visualize the cellular morphology of other pyrenoid-containing algal species such as dinoflagellates^53^ and diatoms^54^. With large-scale fluorescent tagging efforts underway across pyrenoid-containing algae and hornworts to discover new pyrenoid proteins^55,56^, ExM-based pyrenoid protein localization studies similar to this one could accelerate our understanding of pyrenoid organization and biogenesis. They could also generate new hypotheses for protein function and reveal whether similar structural principles are shared across convergently evolved pyrenoids. Finally, combining existing ExM workflows for plant cells^57,58^ with ongoing efforts to engineer pyrenoids into land plants^29,59^ could also enhance the ability to evaluate whether engineered components localize to the correct subdomains. Overall, nanoscale spatial resolution of pyrenoid proteins could help bridge discovery, mechanism, and engineering in the study of pyrenoid biology.

## Acknowledgements

We thank Yana Kazachkova, Ashwani Rai, Micah Burton, Sophie Skanchy and all other members of the Jonikas lab for helpful feedback and discussions. We thank Alistair McCormick for his helpful feedback on the manuscript and Gautam Dey and his lab members for their advice on U-ExM. The BST4 antibody was a generous gift from the Mackinder lab at the University of York, York, UK. Research reported in this publication was supported by the National Institute of General Medical Sciences (NIGMS) of the National Institutes of Health under grant number T32GM007388; by the National Science Foundation under NSF MCB-2410354; by the Bill and Melinda Gates Foundation and United Kingdom Foreign, Commonwealth & Development Office grant INV-054558; and by the Howard Hughes Medical Institute. H.W. is a recipient of a fellowship from the Chinese Research Council. We acknowledge the use of the Imaging and Analysis Center (IAC) operated by the Princeton Materials Institute at Princeton University, which is supported in part by the Princeton Center for Complex Materials (PCCM), a National Science Foundation (NSF) Materials Research Science and Engineering Center (MRSEC; DMR-2011750). The content is solely the responsibility of the authors and does not necessarily represent the official views of the funders.

---

**Figure S1:**
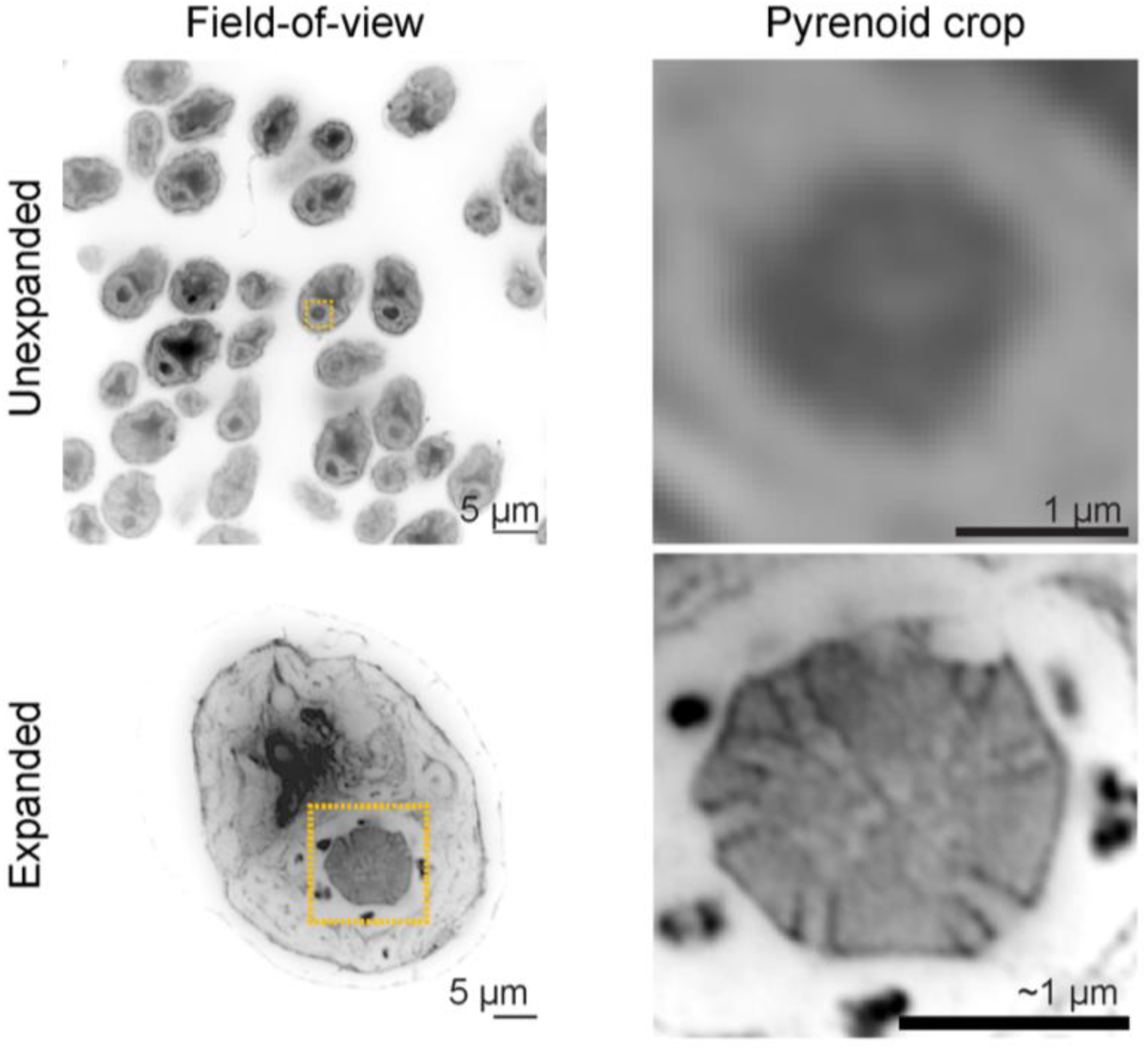
Additional examples of unexpanded and expanded Chlamydomonas cells grown for 6 hours at low CO_2_.

**Figure S2:**
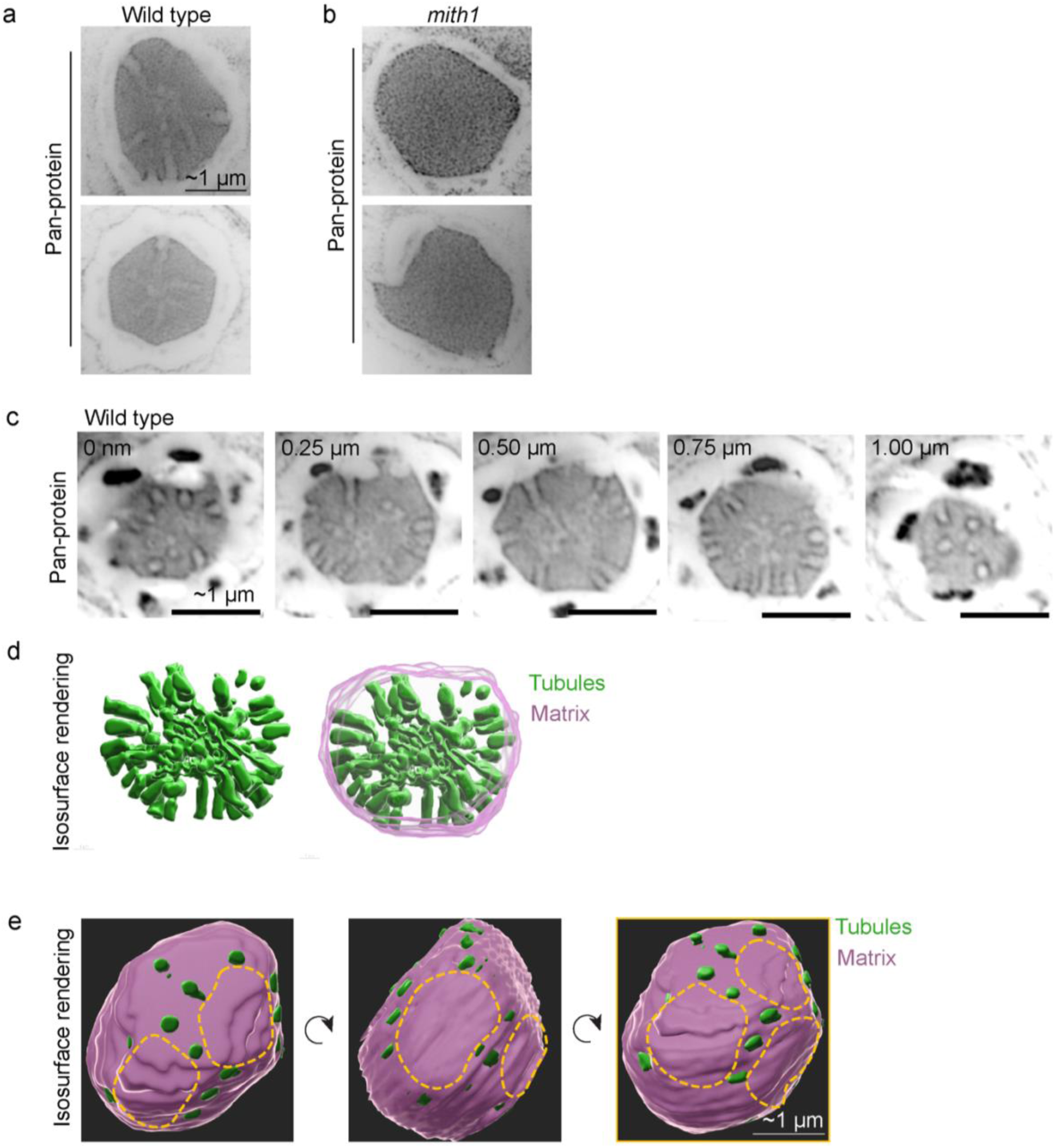
Additional examples of wild-type and *mith1* mutant pyrenoids and wild-type pyrenoid space-filling renderings. **a.** Additional examples of medial slices of wild-type pyrenoids stained with pan-protein stain. **b.** Additional examples of medial slices of *mith1* pyrenoids stained with pan-protein stain. **c.** Confocal z-series of the wild-type pyrenoid from **Figure S1** stained with pan-protein stain. d. Space-filling reconstructions of tubules and matrix segmented from the confocal sections in **c**. **e.** Oblique views of the space-filling rendering in Figure 2c, with the dotted lines representing possible locations of starch granules between the linear arrangements of tubules. Scale bars corrected for expansion factor.

**Figure S3:**
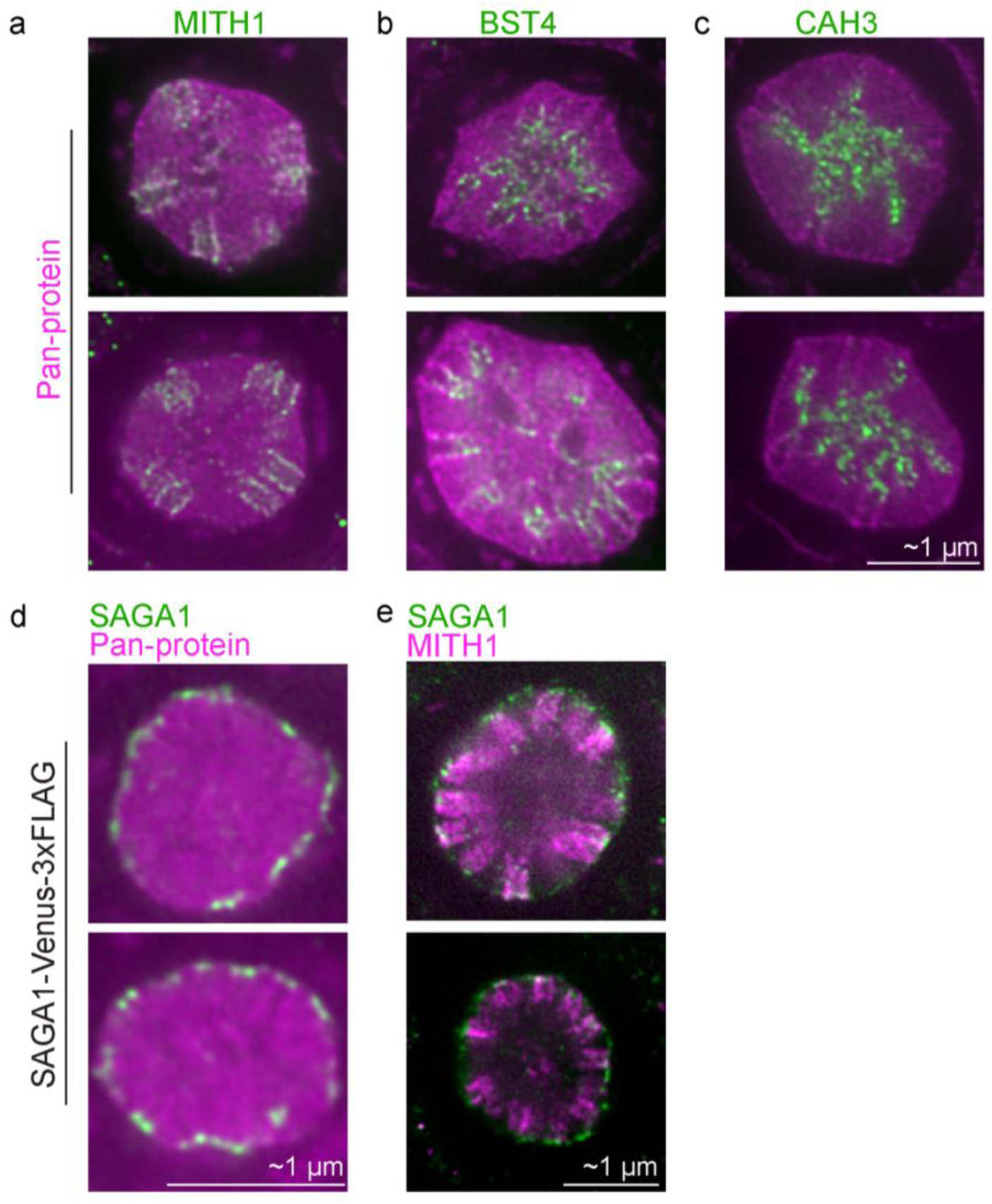
Additional examples of MITH1, BST4, CAH3 and SAGA1 immunofluorescence localization. **a-c.** Wild-type pyrenoids displayed with immunofluorescence (green) merged with pan-protein stain (magenta). The antibodies used were a. MITH1, b. BST4, and c. CAH3. **d-e.** SAGA1-Venus-3xFLAG expressing pyrenoids displayed with **d.** FLAG (green) and pan-protein stain (magenta) and **e.** FLAG (green) and MITH1 (magenta). Medial slices are displayed for each pyrenoid. Scale bars are corrected for expansion factor.

**Figure S4:**
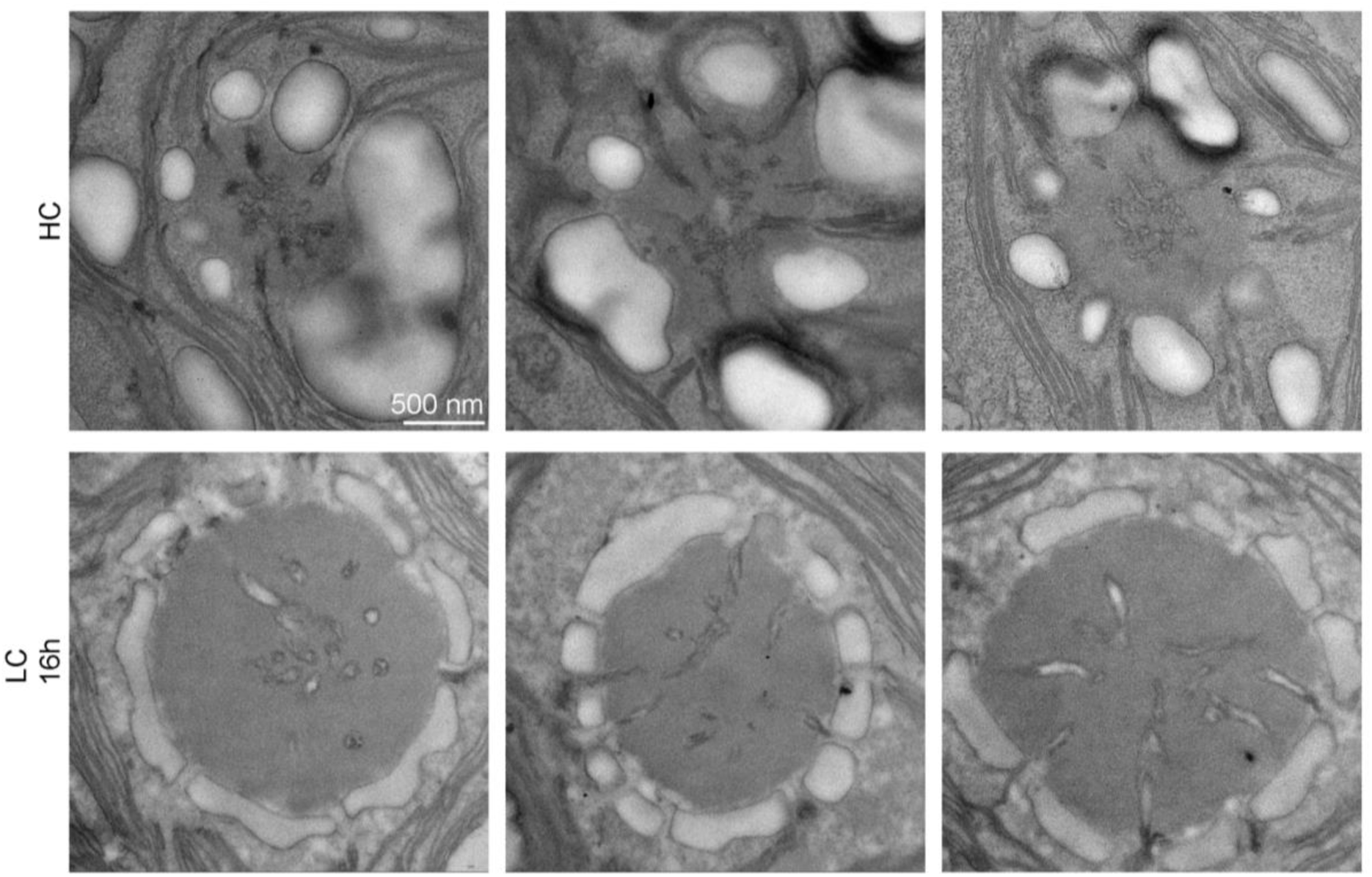
Additional examples of transmission electron micrographs of pyrenoids at high CO_2_ and low CO_2_. Transmission electron micrographs (TEM) of wild-type pyrenoids from cells grown at HC and LC 16h. Scale bar 500 nm.

**Movie S1: U-ExM + iSIM confocal Z-series of a wild-type pyrenoid stained with pan-protein stain**

Scale bar ∼1 µm, corrected for expansion factor. Slices every ∼33 nm, corrected for expansion factor.

**Movie S2: Space-filling rendering of a wild-type pyrenoid matrix (gray) and tubule network (green).**

**Movie S3: Montage of confocal Z-series of wild-type pyrenoids combining immunofluorescence (green) with pan-protein stain (magenta).**

The antibodies used were **a.** MITH1, **b.** BST4 and **c.** CAH3. Scale bars ∼1 µm, expansion corrected.

**Movie S4: Confocal Z-series of the pyrenoid of SAGA1-Venus-3xFLAG-expressing cell combining anti-FLAG (green) and anti-MITH1 (magenta) immunofluorescence.** Scale bar ∼1 µm, expansion corrected.

**Table S1:** List of Chlamydomonas strains used in this study along with their sources.

| Chlamydomonas Resource Center ID | Strain description | Source | Antibiotic resistance |
| --- | --- | --- | --- |
| CC-4533 | CMJ030 | Wild type and parent strain to CLiP1 library | none |
| LMJ.RY0402.133670 | *mith1* | CLiP1 library<br>LMJ.RY0402.133670 | Paromomycin |
| CC-5422 | *saga1*; SAGA1-Venus-3xFLAG | Previously published<sup>46</sup> | Paromomycin,<br>Hygromycin |

## MATERIALS AND METHODS

### Chlamydomonas culture conditions

The strains used in this study can be found in Table S1. All strains were maintained on 1.5% agar containing Tris-acetate phosphate (TAP) medium with revised trace minerals at room temperature (∼22 °C) under very low light (<10 µmol photons⋅m^-2^⋅s^-1^). For liquid culture for U-ExM experiments, strains were inoculated in ∼10 mL of Tris phosphate (TP) medium in 125 mL flasks capped with sterile vented tissue sealing film (MSE supplies, 30 mm vented membrane, 0.2–0.3 µm pore size) while stirring at 180 rpm for aeration under continuous cool white light at ∼150 µmol photons⋅m^-2^⋅s^-1^. For growth in high CO_2_ conditions, strains were cultured in 3% (v/v) CO_2_ for two days before harvesting. For growth in low CO_2_ conditions, strains were first cultured at high CO_2_ as described and then diluted into fresh TP medium before moving to a different incubator with the same light, temperature and stirring parameters except low CO_2_ levels (0.04% v/v) for the indicated amount of time before harvesting. Prior to harvesting, cell counts were measured by a Countess II Automated Cell Counter (Life Technologies).

### Ultrastructure expansion microscopy

Ultrastructure expansion microscopy was performed as described previously^32^ with minor changes. Briefly, 10 mL of Chlamydomonas cells were harvested at the specified timepoint by centrifugation at 600 x g for five minutes and then resuspended in 1 mL of fresh TP medium. 200 µL of each cell suspension was pipetted onto an 18 mm poly-D-lysine coated coverslip (Neuvitro) and 100 µL of the same suspension was pipetted onto a 12 mm poly-D-lysine coated coverslip to be used as the un-expanded control. All coverslips were placed in a 12-well plate and cells were allowed to adhere for ten minutes under ∼120 µmol photons⋅m^-2^⋅s^-1^ white light.

To fix cells, TP was pipetted off the coverslip and 1 mL of ice-cold fixative buffer (4% paraformaldehyde, 0.1% glutaraldehyde in 1X PBS) was added per coverslip and the plate was placed on ice for slow fixation over 60 minutes. Next, the fixative solution was removed and replaced with 1 mL of ice-cold 100 mM glycine solution to quench the fixation for five minutes on ice. The glycine solution was removed, and the coverslips were washed three times in ice-cold 1X PBS. Finally, 1 mL of ice-cold anchoring solution (2% acrylamide, 1.4% formaldehyde in 100 mM bicarbonate buffer) was added to each coverslip and the samples were allowed to anchor at 4 °C overnight with no shaking.

The following day, the gelation chambers were pre-cooled on ice to slow down the gelation. Gelation chambers consisted of glass slides with two 13 mm diameter x 0.6 mm depth silicone spacers adhered to it, placed in a petri dish with damp Whatman paper to retain humidity. The N,N’-methylenebisacrylamide concentration in the monomer solution was reduced following results from the TREx protocol^60^ from 0.1% in the original U-ExM protocol to 0.03% in this work to achieve a higher expansion factor than the U-ExM monomer solution. A freshly thawed 1 mL tube of monomer solution (10% (w/v) acrylamide, 19% (w/v) sodium acrylate, 0.03% (w/v) N,N’-methylenebisacrylamide in 1X PBS) was activated with 25 µL of 10% TEMED and 25 µL of 10% APS and vortexed briefly to mix. 100 µL of activated monomer solution was pipetted into the middle of the silicone spacer of each gelation chamber and the coverslips were removed from anchoring solution, blotted well and inverted cell side down to form the lid of the gelation chamber. Gelation was allowed to proceed for 1 hour on ice. Afterwards, the gels were removed from the gelation chambers and incubated in 6-well plates with 2 mL of denaturation buffer (200 mM SDS, 200 mM NaCl, 50 mM Tris in ddH_2_O, pH 9.0) for 10 minutes with shaking. Once the gels had approximately doubled in size, each gel was cut in half and placed in a 1.5 mL tube filled with fresh denaturation buffer and incubated at 95 °C for 90 minutes. Gels were then transferred back to the 6-well plate for three 1X PBS washes before proceeding to pan-protein staining or immunostaining.

Due to the large size of the gels and the density of cells adhered on the coverslips, each gel could be cut into 6-8 smaller pieces for parallel pan-protein or immunostaining treatments. For pan-protein staining alone, cells were incubated in 20 µg/mL of Atto 565 NHS-ester stain in 100 mM bicarbonate buffer for three hours with shaking at room temperature before the stain was washed out three times for 30 minutes with shaking. For immunostaining alone, cells were incubated with primary antibodies in antibody solution (2% BSA in PBS) overnight at room temperature with shaking. The gels were then washed three times for 10 minutes each with 0.1% PBS-Tween 20 at room temperature before being incubated with secondary antibodies in antibody solution for three hours at room temperature with shaking. The secondary antibody was washed out three times for 10 minutes each with 0.1% PBS-Tween 20 at room temperature. The MITH1^29^, BST4^39^, and CAH3^29^ primary antibodies have been published and validated previously; the FLAG primary antibodies were purchased commercially from Cell Signaling (Rabbit anti-DYKDDDDK Tag Cat No. 14793S, Mouse anti-DYKDDDDK Tag Cat. No. 8146S). All secondary antibodies used were F(ab’)2 fragments from Invitrogen (anti-Rabbit Alexa Fluor 488 Cat No. A-11070, anti-Rabbit Alexa Fluor 564 Cat No. A-11071 and anti-Mouse Alexa Fluor 488 Cat No. A-11017). All primary and secondary antibody dilutions were 1:500 except for primary Mouse anti-FLAG which was diluted to 1:250. For maximum antibody signal retention, the gels were briefly incubated in post-immunostaining fixative solution (0.5% paraformaldehyde in 1X PBS) for five minutes at room temperature with shaking before being washed three times for 10 minutes in 1X PBS. For co-localization between immunofluorescence and pan-protein stain, immunostaining was performed first followed by pan-protein staining. Pre-expansion samples were stained with the same concentration of NHS-ester Atto 565 for the same amount of time as above and washed three times in 1X PBS before mounting.

To expand the samples, gels were placed in 50 mL falcon tubes that were filled with MilliQ water. The water was exchanged twice and then samples were allowed to expand completely overnight before imaging the following day.

### iSIM imaging and image alignment

All light microscopy images were acquired on a VT-iSIM super-resolution spinning disk confocal microscope equipped with a Hamamatsu Orca Quest sCMOS camera and running VisiView control software. Samples were imaged with a 100x 1.35 NA Olympus silicone immersion objective (UPLSAPO100XS) with a z-step of 0.2 µm. For un-expanded images, 5 µL of 100% glycerol was pipetted onto a glass slide and the coverslip containing the cells was inverted onto it. The edges of the coverslip were sealed with clear nail polish before imaging. Two to three fields of view were imaged for each un-expanded sample depending on the cell density. For expanded samples, gels were screened on the microscope to determine which side contained the cells. Then, the gels were placed cell side down onto a 22 mm poly-D-lysine coated coverslip placed in an Attofluor cell chamber. ∼1 mL of warm 1% low melting point agarose in water was flowed over the gel to prevent desiccation during long imaging sessions. Atto 565 NHS-ester and Alexa Fluor 568-conjugated secondary antibody were excited with a 561 nm laser and collected at 595/40 nm. Alexa Fluor 488-conjugated secondary antibody was excited with a 488 nm laser and collected at 525/25 nm. At least seven cells were imaged per expanded sample.

All images were deconvolved using the Microvolution plugin for Fiji^61^. To correct for misalignment between imaging channels, 100 nm Tetraspeck beads were imaged with both imaging channels and the images were deconvolved. The offset between the channels was determined from the images of the beads and corrected manually using Fiji. The same offset was then applied to all two-channel ExM images to correct for system misalignment.

### Transmission electron microscopy

Transmission electron microscopy was performed as described previously^62^ with some minor changes. For cells grown at high CO_2_, cells were cultured with stirring as described above. For cells grown at low CO_2_, cells were cultured in TAP medium under ambient (0.04% v/v) CO_2_, before being pelleted and resuspended in TP at ambient CO_2_ for 16 hours prior to being harvested. All cultures were harvested at a cell density of 1-2 x 10^6^ cells mL^-1^. At least 5 x 10^7^ cells were collected by centrifugation at 600 x g for five minutes in 50 mL tubes, before proceeding with fixation, staining, resin embedding and sectioning as previously described. All imaging was performed using a Talos L120C G2 (S)TEM (Thermo Fisher Scientific) at the Imaging and Analysis Center, Princeton University.

### Expansion factor and circularity measurements

While the distortion in ExM protocols is generally low^63^, we reasoned that the protein-dense phase-separated pyrenoid could have a different expansion factor than the whole cell. To account for this possible variability in expansion, we calculated the expansion factor of each experiment based on the diameter of the pyrenoid before and after expansion. For every genotype or growth condition, the pyrenoid was outlined with the free-hand selection tool in Fiji in un-expanded and expanded cells. The major axis, minor axis and circularity were measured for each free-hand selection. Expansion factor was calculated as the ratio between the average expanded major axis measurement and the average un-expanded major axis measurement. Circularity was calculated by Fiji as 4π x (area/perimeter^2^).

### Immunofluorescence distribution analysis

To analyze the distribution of immunofluorescence signal across the axis of the pyrenoid in Fiji, the pan-protein staining channel was used to outline the pyrenoid using the free-hand selection tool and the selection was added to the ROI manager. For images where the pan-protein staining channel was unavailable, the most peripheral protein signal was used to draw the free-hand selection. Fluorescence intensity as a function of radial distance from the pyrenoid centroid was calculated for each channel using the Radial Profile Plot plugin (https://imagej.net/ij/plugins/radial-profile.html). Briefly, the plugin fits a circle to the free-hand selection and generates a plot of fluorescence intensity within concentric circles as a function of distance from the center of the fitted circle. The data were normalized in two ways. To allow pyrenoids of different sizes to be averaged for comparison, all radial distance values were normalized by dividing by the radius of the fitted circle, resulting in normalized radial distances ranging from 0 to 1. Fluorescence intensity measurements for each channel were first background corrected and then normalized to the maximum pixel intensity within that channel, resulting in relative fluorescence intensity values ranging from 0 to 1. The fluorescence intensities were binned using a bin size of 0.05 normalized radial units to reduce unequal weighting of larger pyrenoids, which contained more radial data points than smaller pyrenoids.

To analyze the distribution of immunofluorescence signal across the transverse axis of individual tubules in Fiji, the straight-line tool with a width of 5 pixels was used to draw lines across two tubules per pyrenoid using the pan-protein channel: one cross-section view from a superficial slice and one longitudinal section view from a medial slice of a z-stack. The lines were added to the ROI manager and the Plot Profile tool in Fiji was used to measure fluorescence intensity of both channels along the line. The Find Peaks BAR plugin (https://imagej.net/plugins/find-peaks) was used on the pan-protein distribution plot to find the x-axis location of the two pan-protein peaks. The data were normalized in two ways. To align the peaks and troughs of pan-protein staining across multiple tubules, we calculated a normalized position across the tubule axis (x_norm_) such that the trough at the midpoint between the peaks was aligned at 0 and the left and right peaks were positioned at approximately -1 and 1, respectively. The positions of the left (x_left_) and right peaks (x_right_) were used to find the position of the center of the peaks X_center_ = ((x_left_ + x_right_)/2) and the halfwidth of the tubule x_hw_ = ((x_right_ - x_left)_/2). The normalized x position of each point (x_norm_) was calculated as (x – x_center_) / x_hw_. The fluorescence intensity measurements for each channel were background corrected and then normalized to the maximum pixel intensity within that channel, resulting in relative fluorescence intensity values ranging from 0 to 1. The fluorescence intensities were binned using a bin size of 0.05 normalized positional units to reduce unequal weighting of wider peripheral tubules, which contained more data points than narrower medial tubules.

### Measuring tubule diameter in U-ExM and TEM images

To measure the tubule diameter from U-ExM images, the straight-line tool with a width of 5 pixels was used to draw lines across two tubules per pyrenoid using the pan-protein channel – one cross-section view from a superficial slice and one peripheral longitudinal section view from a medial slice of a z-stack. The Plot Profile tool was used to measure the fluorescent intensity distribution along the line and then the Find Peaks BAR plugin was used on the pan-protein distribution plot to find the x-axis location of the two pan-protein peaks (x_right_ and x_left_). The tubule diameter was then calculated as x_right_ - x_left_ and divided by the expansion factor of that sample to get the expansion corrected tubule diameter.

Our TEM protocol is optimized to achieve high contrast when looking at membranes. To measure the tubule diameter from TEM images, the straight-line tool in Fiji was used to draw a line that measured from one side of the electron dense membrane to the other. Only longitudinal sections of the tubules were measured at the periphery of the pyrenoid matrix, and two tubules were measured per micrograph when possible.

### Space-filling rendering of matrix and tubules

Volumetric isosurface renderings of the wild-type pyrenoid matrix and tubules were achieved using Imaris (Version 11.0.0). The pyrenoid matrix was thresholded automatically using the Surfaces tool. Due to the tubules appearing as signal-sparse structures within the matrix, they had to be manually segmented in every slice using the “Magic Wand” selection tool within the Surfaces tool.

### Statistical analysis

All statistical analyses were performed in R. Significance between two groups was determined using a two-tailed t-test. For comparisons of three or more groups, we used a Kruskal-Wallis rank sum test followed by a post-hoc Dunn’s test with a Bonferroni correction for multiple comparisons.

## REFERENCES

1. He, S., Crans, V. L. & Jonikas, M. C. The pyrenoid: the eukaryotic CO2-concentrating organelle. Plant Cell 35, 3236–3259 (2023).

2. Field, C. B., Behrenfeld, M. J., Randerson, J. T. & Falkowski, P. Primary Production of the Biosphere: Integrating Terrestrial and Oceanic Components. Science 281, 237–240 (1998).

3. Barrett, J., Nam, O., Naduthodi, M. I. S. & Mackinder, L. C. M. Pyrenoid Structure, Function, Evolution, and Characterization Across Diverse Lineages. Annu. Rev. Plant Biol. 77, 81–115 (2026).

4. Mackinder, L. C. M. et al. A repeat protein links Rubisco to form the eukaryotic carbon-concentrating organelle. Proc. Natl. Acad. Sci. U. S. A. 113, 5958–5963 (2016).

5. Mackinder, L. C. M. et al. A Spatial Interactome Reveals the Protein Organization of the Algal CO2-Concentrating Mechanism. Cell 171, 133–147.e14 (2017).

6. Wang, L. et al. A chloroplast protein atlas reveals punctate structures and spatial organization of biosynthetic pathways. Cell 186, 3499–3518.e14 (2023).

7. Freeman Rosenzweig, E. S., et al. The Eukaryotic CO2-Concentrating Organelle Is Liquid-like and Exhibits Dynamic Reorganization. Cell 171, 148–162.e19 (2017).

8. He, S. et al. The structural basis of Rubisco phase separation in the pyrenoid. Nat. Plants 6, 1480–1490 (2020).

9. Wunder, T., Cheng, S. L. H., Lai, S.-K., Li, H.-Y. & Mueller-Cajar, O. The phase separation underlying the pyrenoid-based microalgal Rubisco supercharger. Nat. Commun. 9, 5076 (2018).

10. Engel, B. D. et al. Native architecture of the Chlamydomonas chloroplast revealed by in situ cryo-electron tomography. eLife 4, e04889 (2015).

11. Ohad, I., Siekevitz, P. & Palade, G. E. Biogenesis of chloroplast membranes. II. Plastid differentiation during greening of a dark-grown algal mutant (Chlamydomonas reinhardi). J. Cell Biol. 35, 553–584 (1967).

12. Sager, R. & Palade, G. E. Structure and development of the chloroplast in Chlamydomonas : I The normal green cell. J. Biophys. Biochem. Cytol. 3, 463–488 (1957).

13. Badger, M. R. et al. The diversity and coevolution of Rubisco, plastids, pyrenoids, and chloroplast-based CO2-concentrating mechanisms in algae. Can. J. Bot. 76, 1052–1071 (1998).

14. Hanson, D. T., Franklin, L. A., Samuelsson, G. & Badger, M. R. The Chlamydomonas reinhardtii cia3 Mutant Lacking a Thylakoid Lumen-Localized Carbonic Anhydrase Is Limited by CO2 Supply to Rubisco and Not Photosystem II Function in Vivo. Plant Physiol. 132, 2267–2275 (2003).

15. Raven, J. A. CO2-concentrating mechanisms: a direct role for thylakoid lumen acidification? Plant Cell Environ. 20, 147–154 (1997).

16. Fei, C., Wilson, A. T., Mangan, N. M., Wingreen, N. S. & Jonikas, M. C. Modelling the pyrenoid-based CO2-concentrating mechanism provides insights into its operating principles and a roadmap for its engineering into crops. Nat. Plants 8, 583–595 (2022).

17. Toyokawa, C., Yamano, T. & Fukuzawa, H. Pyrenoid Starch Sheath Is Required for LCIB Localization and the CO2-Concentrating Mechanism in Green Algae. Plant Physiol. 182, 1883–1893 (2020).

18. Karlsson, J. et al. A novel alpha-type carbonic anhydrase associated with the thylakoid membrane in Chlamydomonas reinhardtii is required for growth at ambient CO2. EMBO J. 17, 1208–1216 (1998).

19. Karlsson, J., Hiltonen, T., Husic, H. D., Ramazanov, Z. & Samuelsson, G. Intracellular Carbonic Anhydrase of Chlamydomonas reinhardtii. Plant Physiol. 109, 533–539 (1995).

20. Moroney, J. V., Tolbert, N. E. & Sears, B. B. Complementation analysis of the inorganic carbon concentrating mechanism of Chlamydomonas reinhardtii. Mol. Gen. Genet. MGG 204, 199–203 (1986).

21. Spalding, M. H., Spreitzer, R. J. & Ogren, W. L. Carbonic Anhydrase-Deficient Mutant of Chlamydomonas reinhardii Requires Elevated Carbon Dioxide Concentration for Photoautotrophic Growth. Plant Physiol. 73, 268–272 (1983).

22. Pronina, N. A. & Semenenko, V. E. The role of pyrenoid in concentrating generation and fixation of CO2 in chloroplasts of microalgae. Role Pyrenoid Conc. Gener. Fixat. CO2 Chloroplasts Microalgae 39, 723–732 (1992).

23. Catherall, E. et al. From algae to plants: understanding pyrenoid-based CO2-concentrating mechanisms. Trends Biochem. Sci. 50, 33–45 (2025).

24. Adler, L. et al. New horizons for building pyrenoid-based CO2-concentrating mechanisms in plants to improve yields. Plant Physiol. 190, 1609–1627 (2022).

25. Long, S. P., Marshall-Colon, A. & Zhu, X.-G. Meeting the global food demand of the future by engineering crop photosynthesis and yield potential. Cell 161, 56–66 (2015).

26. Ohad, I., Siekevitz, P. & Palade, G. E. Biogenesis of chloroplast membranes. I. Plastid dedifferentiation in a dark-grown algal mutant (Chlamydomonas reinhardi). J. Cell Biol. 35, 521–552 (1967).

27. Borkhsenious, O. N., Mason, C. B. & Moroney, J. V. The Intracellular Localization of Ribulose-1,5-Bisphosphate Carboxylase/Oxygenase in Chlamydomonas reinhardtii. Plant Physiol. 116, 1585–1591 (1998).

28. Gustafsson, M. G. L. Surpassing the lateral resolution limit by a factor of two using structured illumination microscopy. J. Microsc. 198, 82–87 (2000).

29. Hennacy, J. H. et al. SAGA1 and MITH1 produce matrix-traversing membranes in the CO2-fixing pyrenoid. Nat. Plants 10, 2038–2051 (2024).

30. Hell, S. W. & Wichmann, J. Breaking the diffraction resolution limit by stimulated emission: stimulated-emission-depletion fluorescence microscopy. Opt. Lett. 19, 780–782 (1994).

31. Curd, A. et al. Construction of an instant structured illumination microscope. Methods 88, 37–47 (2015).

32. Gambarotto, D., Hamel, V. & Guichard, P. Ultrastructure expansion microscopy (U-ExM). Methods Cell Biol. 161, 57–81 (2021).

33. Klena, N. et al. An In-depth Guide to the Ultrastructural Expansion Microscopy (U-ExM) of *Chlamydomonas reinhardtii*. Bio-Protoc. 13, (2023).

34. Laporte, M. H., Klena, N., Hamel, V. & Guichard, P. Visualizing the native cellular organization by coupling cryofixation with expansion microscopy (Cryo-ExM). Nat. Methods 19, 216–222 (2022).

35. Yu, C.-C. (Jay) et al. Expansion microscopy of C. elegans. eLife 9, e46249 (2020).

36. Mao, C. et al. Feature-rich covalent stains for super-resolution and cleared tissue fluorescence microscopy. Sci. Adv. 6, eaba4542 (2020).

37. M’Saad, O. & Bewersdorf, J. Light microscopy of proteins in their ultrastructural context. Nat. Commun. 11, 3850 (2020).

38. Meyer, M. T. et al. Assembly of the algal CO2-fixing organelle, the pyrenoid, is guided by a Rubisco-binding motif. Sci. Adv. 6, eabd2408 (2020).

39. Adler, L. et al. The role of BST4 in the pyrenoid of Chlamydomonas reinhardtii. 2023.06.15.545204 Preprint at 10.1101/2023.06.15.545204 (2023).

40. Ergun, S. L., et al. A bifunctional coiled-coil protein generates the membrane-within-condensate architecture of the CO₂-fixing pyrenoid. Submitt. Biorxiv (2026).

41. Blanco-Rivero, A., Shutova, T., Román, M. J., Villarejo, A. & Martinez, F. Phosphorylation Controls the Localization and Activation of the Lumenal Carbonic Anhydrase in Chlamydomonas reinhardtii. PLoS ONE 7, e49063 (2012).

42. Markelova, A. G., Sinetova, M. P., Kupriyanova, E. V. & Pronina, N. A. Distribution and functional role of carbonic anhydrase Cah3 associated with thylakoid membranes in the chloroplast and pyrenoid of Chlamydomonas reinhardtii. Russ. J. Plant Physiol. 56, 761–768 (2009).

43. Mitra, M. et al. The carbonic anhydrase gene families of Chlamydomonas reinhardtii. Can. J. Bot. 83, 780–795 (2005).

44. Park, Y.-I. et al. Role of a novel photosystem II-associated carbonic anhydrase in photosynthetic carbon assimilation in Chlamydomonas reinhardtii. FEBS Lett. 444, 102–105 (1999).

45. Wang, L. et al. Chloroplast-mediated regulation of CO2-concentrating mechanism by Ca2+-binding protein CAS in the green alga Chlamydomonas reinhardtii. Proc. Natl. Acad. Sci. 113, 12586–12591 (2016).

46. Itakura, A. K. et al. A Rubisco-binding protein is required for normal pyrenoid number and starch sheath morphology in Chlamydomonas reinhardtii. Proc. Natl. Acad. Sci. U. S. A. 116, 18445–18454 (2019).

47. Crans, V. L., Burton, M. I., Garde, A., Wang, L. & Jonikas, M. C. SAGA1 and SAGA2 localize the starch sheath to the pyrenoid in Chlamydomonas reinhardtii. Proc. Natl. Acad. Sci. 123, e2533609123 (2026).

48. Ramazanov, Z. et al. The induction of the CO2-concentrating mechanism is correlated with the formation of the starch sheath around the pyrenoid of Chlamydomonas reinhardtii. Planta 195, 210–216 (1994).

49. Ma, Y., Pollock, S. V., Xiao, Y., Cunnusamy, K. & Moroney, J. V. Identification of a Novel Gene, CIA6, Required for Normal Pyrenoid Formation in Chlamydomonas reinhardtii. Plant Physiol. 156, 884–896 (2011).

50. Caspari, O. D. et al. Pyrenoid loss in Chlamydomonas reinhardtii causes limitations in CO2 supply, but not thylakoid operating efficiency. J. Exp. Bot. 68, 3903–3913 (2017).

51. Benlloch, R. et al. Crystal Structure and Functional Characterization of Photosystem II-Associated Carbonic Anhydrase CAH3 in Chlamydomonas reinhardtii1[OPEN]. Plant Physiol. 167, 950–962 (2015).

52. Moroney, J. V., Bartlett, S. G. & Samuelsson, G. Carbonic anhydrases in plants and algae. Plant Cell Environ. 24, 141–153 (2001).

53. Rao, A. K. et al. Hijacking and integration of algal plastids and mitochondria in a polar planktonic host. Curr. Biol. 35, 2509–2523.e7 (2025).

54. Flori, S. et al. Diatom ultrastructural diversity across controlled and natural environments. Curr. Biol. 35, 5709–5720.e4 (2025).

55. Nam, O. et al. A protein blueprint of the diatom CO2-fixing organelle. Cell 187, 5935–5950.e18 (2024).

56. Robison, T. A. et al. Hornworts reveal a spatial model for pyrenoid-based CO2-concentrating mechanisms in land plants. Nat. Plants 11, 63–73 (2025).

57. Cox Jr., K. L., et al. ExPOSE: a comprehensive toolkit to perform expansion microscopy in plant protoplast systems. Plant J. 121, e70049 (2025).

58. Hawkins, T. J., Robson, J. L., Cole, B. & Bush, S. J. Expansion Microscopy of Plant Cells (PlantExM). in The Plant Cytoskeleton: Methods and Protocols (eds Hussey, P. J. & Wang, P.) 127–142 (Springer US, New York, NY, 2023). doi:10.1007/978-1-0716-2867-6_10.

59. Robison, T. A. et al. An unconventional Rubisco small subunit underpins the CO2-concentrating organelle in land plants. Science 391, 1070–1075 (2026).

60. Damstra, H. G. et al. Visualizing cellular and tissue ultrastructure using Ten-fold Robust Expansion Microscopy (TREx). eLife 11, e73775 (2022).

61. Schindelin, J., et al. Fiji: an open-source platform for biological-image analysis. Nat. Methods 9, 676–682 (2012).

62. Franklin, E. et al. Proteomic analysis of the pyrenoid-traversing membranes of Chlamydomonas reinhardtii reveals novel components. New Phytol. 249, 359–372 (2026).

63. Gaudreau-Lapierre, A., Mulatz, K., Béïque, J.-C. & Trinkle-Mulcahy, L. Expansion microscopy-based imaging of nuclear structures in cultured cells. STAR Protoc. 2, 100630 (2021).

